# Brr2 is a splicing fidelity factor

**DOI:** 10.1101/354514

**Authors:** Megan Mayerle, Christine Guthrie

## Abstract

Many spliceosomal DExD/H box helicases act as fidelity factors during pre-mRNA splicing, promoting on-pathway interactions while simultaneously minimizing errors. Mutations linked to Retinitis Pigmentosa (RP), a form of heritable blindness, map to key domains of spliceosomal helicase Brr2 (*SNRNP200* in humans). Previous data show that such mutations negatively impact spliceosome activation, likely due to defects in *brr2-RP* RNA binding, helicase, and ATPase activities. Furthermore, data from human reporter constructs suggest that *brr2-RP* might impact 5′ splice site selection. Here we undertake a systematic analysis of *brr2-RP* effects on splicing fidelity. We show that a subset of *brr2-RP* mutants exhibit intron retention *in vivo*. Furthermore, *brr2-RP* mutants display hyperaccurate and/or error-prone splicing of a variety of splicing reporters. Branch-site fidelity is particularly impacted in this reporter assay. In addition, multiple *brr2-RP* alleles genetically interact with *prp16* alleles known to impact the fidelity of branch site selection. Together these data implicate Brr2 in the fidelity of branch-site selection, and suggest that RP results not just from defects in spliceosome activation, but also from fidelity defects arising throughout the splicing cycle and in splicing fidelity.

## Introduction

Splicing is an essential step in gene expression wherein the spliceosome identifies and catalyzes the removal of non-coding introns and the joining of flanking exons from pre-mRNAs (Will and Lührmann 2011). Splicing is an inherently high-fidelity reaction, with an error rate as low as 1 in every 100,000 splicing events (Fox-Walsh and Hertel 2009). This high level of fidelity results from several independent quality-control pathways as well as from fidelity mechanisms enacted by the spliceosome’s own RNA and protein components, particularly the DExD/H box ATPases. Spliceosomal DExD/H box ATPases, such as Prp5, Prp16, and Prp22, employ kinetic proofreading to both antagonize the splicing of suboptimal, low fidelity substrates and to promote the splicing of optimal, high fidelity substrates (Semlow and Staley 2012).

Spliceosomal protein Brr2 is a DEIH box ATPase and the most highly processive spliceosomal helicase(Raghunathan and Guthrie 1998). Initial Brr2 characterizations focused on its role in unwinding the snRNA duplexes during spliceosome activation (Lauber et al. 1996; Raghunathan and Guthrie 1998), however Brr2 has since been implicated in multiple steps of the splicing cycle: during the transition between the 1^st^ and 2^nd^ catalytic steps (Hahn et al. 2012; Mayerle and Guthrie 2016), during spliceosome disassembly (Small et al. 2006), and in pre-mRNA discard from the spliceosome (Chen et al. 2013). Brr2’s helicase and ATPase activities are highly regulated, via both intramolecular and extra-molecular mechanisms, to ensure that Brr2 acts only at appropriate points in the splicing cycle(Absmeier et al. 2017).

Brr2 is one of multiple spliceosomal proteins implicated in *Retinitis Pigmentosa* (RP), a hereditary form of blindness(Růžičková and Staněk 2017). Point mutations identified as causal in RP tend to map to highly conserved, and thus likely functionally important, amino acids. Numerous groups have created yeast strains wherein the genes coding for splicing factors have homologous mutations to those implicated in RP, and used such yeast to probe spliceosome assembly and function (reviewed in (Růžičková and Staněk 2017; Daiger et al. 2014).

Here we show that *brr2-RP* yeast impact a wide variety of Brr2 functions in the cell. Most importantly, we show that *brr2-RP* affect splicing efficiency and fidelity, exhibiting both error-prone and hyperaccurate splicing depending upon the pre-mRNA substrate. We also genetically link Brr2 and Prp16 ATPase activities. While the underlying mechanism remains unclear, together our data clearly implicate Brr2 in splicing fidelity, and suggest that splicing fidelity, not just splicing efficiency, may be impacted in RP.

## Results and Discussion

### Alleles of *BRR2* linked to retinitis pigmentosa impact multiple Brr2 functions both *in vitro* and *in vivo*

We created *S. cerevesiae* strains bearing alleles of *BRR2* that are homologous to specific alleles of *SNRNP200* implicated in RP pathogenesis in humans(Růžičková and Staněk 2017). Working in yeast allows us to take advantage of the extensive array of genetic, biochemical, and molecular techniques available within this system to provide mechanistic insights into how *brr2-RP* impact Brr2 function. *brr2-RP* alleles *brr2-C520R* and *brr2-Q904E* both map to Brr2’s ATPase domain, while alleles *N1104A, R1107A*, and *R1107L* map to the ratchet helix in Brr2’s helicase domain, both of which are essential for Brr2 function (Figure 1A)(Santos et al. 2012; Nguyen et al. 2013). The *brr2-R1107M* allele included in our studies has not been associated with RP; it was a cloning error that serves as a control. In all of these strains, the genomic copy of *BRR2* has been deleted and Brr2 and Brr2-RP variants are expressed, under the control of their endogenous promoter, from a pRS313 plasmid.

Given that Brr2 is an essential protein(Noble and Guthrie 1996; Raghunathan and Guthrie 1998), we initially set out to assess how *brr2-RP* impact yeast growth. We spotted serial dilutions of log-phase yeast cultures onto rich media and allowed *BRR2* and *brr2-RP* strains to grow at 37°C, 30°C, 25°C, and 17°C. Consistent with previous results, *brr2-R1107A* and *brr2-R1107L* exhibited cold sensitive growth at both 25°C and 17°C (Figure 1B,C)(Pena et al. 2009; Zhao et al. 2009; Small et al. 2006; Cordin et al. 2014; Hahn et al. 2012). Previous studies have shown that both Brr2-R1107A and Brr2-R1107L also exhibit decreased helicase activity *in vitro* (Figure 1C, and references indicated in figure legend). However, *brr2-R1107M*, which is not associated with RP, grew as wild type *BRR2*, clearly indicating that not all mutations at position R1107 negatively impact yeast growth (Figure 1B). Furthermore, Brr2-Q904E and Brr2-N1104L also exhibit compromised helicase activity(Zhao et al. 2009; Mozaffari-Jovin et al. 2013; Ledoux and Guthrie 2016), but neither mutation causes cold sensitivity (Figure 1C). While it is possible that helicase activity is not compromised enough by these alleles to cause cold sensitivity, it is also likely that other aspects of Brr2 function may also be negatively affected by Brr2-R1107A and Brr2-R1107L and contribute to the cold sensitive phenotype.

**Figure 1:**
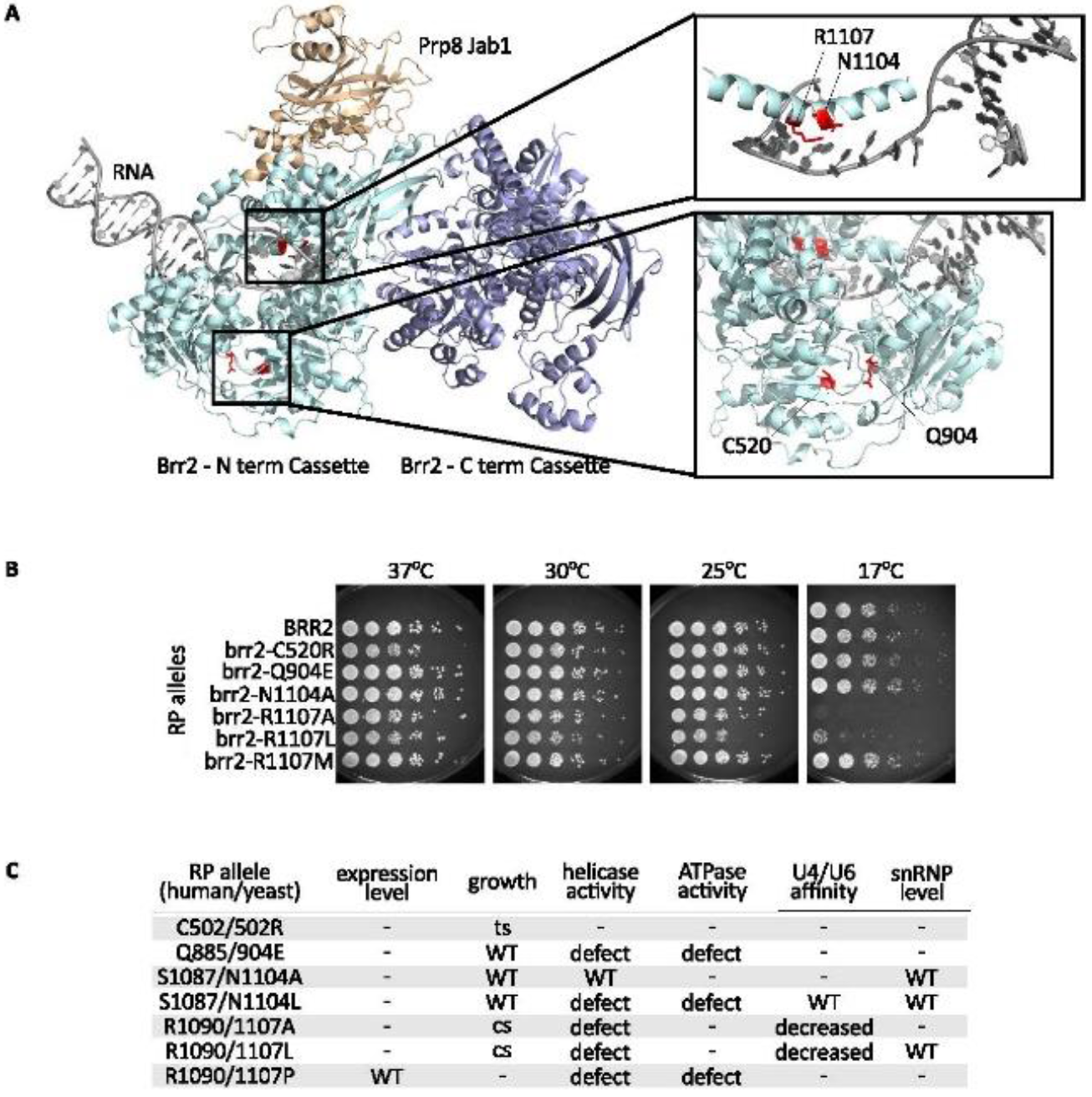
*brr2-RP* impact multiple aspects of Brr2 function during splicing. A. Structure of Brr2′s N (cyan) and C (blue) cassettes(Santos et al. 2012). Amino acid positions impacted in RP are colored in red. The Prp8 Jab1 domain is shown in tan, and a modeled RNA duplex in gray. A-top right. Zoom in on positions C520 and Q904, which are located proximate to Brr2′s ATPase domain. A-bottom right. Zoom in on positions N1104 and R1107 in Brr2′s ratchet helix, which allows for processive dsRNA unwinding by Brr2. Figures made using pymol, PDB:4BGD (www.pymol.org). B. Spotting assay showing growth of different *brr2-RP* on rich media at the temperatures indicated. A subset of *brr2-RP* exhibit cold (R1107A, R1107L) or temperature (C520R) sensitive growth. C. Table summarizing known effects of *brr2-RP*. Data are included from this work, and the following references: (Zhao et al. 2009; Zhang et al. 2009; Santos et al. 2012; Mozaffari-Jovin et al. 2013; Ledoux and Guthrie 2016).

By contrast, *brr2-C520R* was temperature sensitive, exhibiting a slight but repeatable growth defect at 37°C (Figure 1B). To our knowledge, this is the only temperature sensitive *brr2-RP* allele. We speculate that *brr2-C520R* temperature sensitivity could result from a defect in ATP binding caused by destabilization of the Brr2 ATP-binding pocket, however more biochemical assays are required to understand how this defect arises. Together these data show that RP-associated amino acid substitutions negatively impact a wide range of Brr2 functions.

### Multiple endogenous pre-mRNAs exhibit intron retention in *brr2-RP*

The effects of RP mutations on Brr2 growth, ATPase activity, and helicase activity that we and others have observed suggest that pre-mRNA splicing may be compromised in *brr2-RP in vivo*. To test this we measured the relative abundancies of pre-mRNA and spliced mRNA of six transcripts (*ACT1*, *DBP2*, *ERV1*, *NSP1*, *SEC17*, and *UBC5*) by RT-qPCR. Both *ACT1* and *NSP1* encode pre-mRNA transcripts with introns that can be considered typical for *S. cerevisiae* in both size (308 and 118 nt, respectively) and splice-site sequence (Spingola et al. 1999). *DBP2* encodes a pre-mRNA transcript with an unusually long intron (over 1000 nt). *UBC5* and *SEC17* both encode pre-mRNA transcripts with atypical aAG 3′ splice sites. *ERV1* transcripts have a non-consensus UAuUAAC branch site(Engel et al. 2014). In addition, while all of these genes are constitutively expressed, their relative levels of expression vary significantly(Thompson and Cubillos 2017).

**Figure 2:**
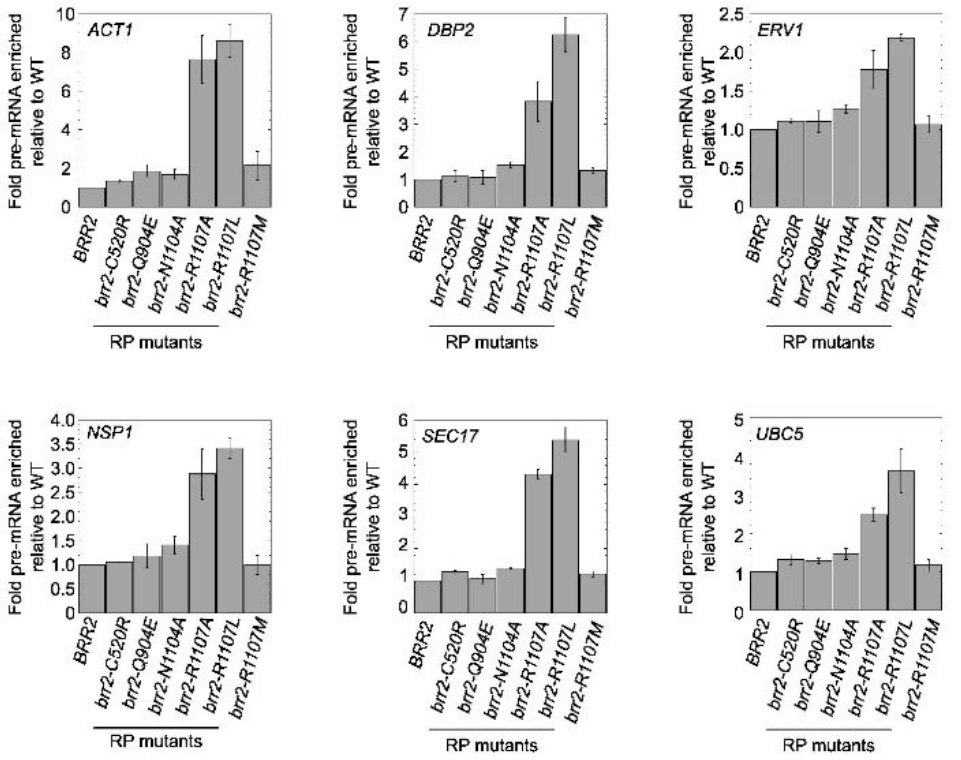
Multiple cellular transcripts exhibit intron retention *in vivo* in *brr2-RP*. RT-qPCR showing level of pre-mRNA accumulation in *brr2-RP. brr2-R1107A* and *brr2-R1107L* consistently exhibit elevated levels of unspliced pre-mRNA relative to wild type *BRR2*. Experiments performed with total RNA isolated from log phase cultures grown in rich media at 30°C.

We performed RT-qPCR on cDNA made from total RNA isolated from log phase wild type *BRR2*, the five *brr2-RP* strains, and also the control *brr2-R1107M* strain grown at 30°C (Figure 2). Regardless of transcript identity, only *brr2-R1107A* and *brr2-R1107L* exhibit significant intron retention consistent with a defect in pre-mRNA splicing (Figure 2). Both alleles are cold sensitive alleles (Figure 1B). However, though *brr2-R1107A* is slightly more cold sensitive than *brr2-R1107L*, in all cases the *brr2-R1107L* intron retention defect is stronger (Figure 1,2). No other *brr2* alleles showed any appreciable intron retention. That there is no intron retention defect in *brr2-R1107M* suggests that, as with cold sensitivity, not all substitutions of amino acid R1107 are detrimental. Furthermore, though helicase activity is compromised in *brr2-Q904E*, this strain does not exhibit cold sensitivity or intron retention (Figure 1B, C and Figure 2), suggesting that defects in helicase activity per se are not the direct or sole cause of intron retention in *brr2-R1107A* and *R1107L*, though it is also possible that the Q904E mutation simply does not have enough of an effect on helicase activity to impact intron retention. More experiments are needed to ascertain what, precisely, causes intron retention in this strain.

### Brr2 is a splicing fidelity factor

Given that multiple spliceosomal DExD/H box proteins use kinetic proofreading mechanisms to ensure that splicing proceeds with high fidelity(Semlow and Staley 2012), and that RP-associated variants of human Brr2 have been linked to cryptic splicing(Cvačková et al. 2013), we were curious as to whether *brr2-RP* impacted splicing fidelity. We decided to use Act-Cup splicing reporters to assess whether or not *brr2-RP* affect splicing fidelity. The Act-Cup system is an established assay that reports on both splicing fidelity and efficiency(Lesser and Guthrie 1993). In this system, the well-characterized *ACT1* intron is fused to the *CUP1* coding sequence (Figure 3B). The Cup1 protein chelates copper in a concentration-dependent manner, allowing yeast to survive in what would otherwise be lethal copper-containing media. By fusing the *ACT1* intron upstream of *CUP1*, this reporter directly links splicing efficiency to Cup1 expression level, which is assessed by comparing growth of *brr2-RP* and *BRR2* yeast strains transformed with the Act-Cup reporter in copper-containing media (Figure 3A). Splicing fidelity can be gauged by mutating the 5′ splice site, branch site, or 3′ splice site of the Act-Cup reporter and comparing the growth of *brr2-RP* to *BRR2* strains (Figure 3C). Increased growth indicates lower fidelity splicing, and decreased growth indicates higher fidelity and/or lower efficiency splicing.

We knocked out the genomic copy of *CUP1* in our *BRR2* and *brr2-RP* strains and transformed them with the wild type Act-Cup reporter as well as a variety of Act-Cup reporters bearing mutations in their 5′ splice site, branch site, and 3′ splice site. Serial dilutions of log phase cultures of these strains were spotted onto plates supplemented with between 0 and 1.5 mM CuSO_4_ and their growth at 25°C, 30°C, and 37°C assessed. We determined the maximal concentration of CuSO_4_ that supported growth for each *brr2-RP* carrying each reporter [Cu_max_, *brr2-RP*]. We compared this value for the maximum concentration of growth tolerated by *BRR2* bearing the same reporter [Cu_max_, *BRR2*], and used them to calculate a splicing index log_2_([Cu_max_, *brr2-RP*]/([Cu_max_, *brr2-RP*]+ [Cu_max_, *BRR2*]). We summarize these data as a heat map in Figure 3D. Blue coloring indicates that the indicated *brr2-RP* Act-Cup reporter combination grew less well than the same reporter in *BRR2*, yellow coloring indicates better growth (Figure 3D). We included *brr2-1* as a non-RP control known to show defects in dsRNA unwinding(Raghunathan and Guthrie 1998; Maeder et al. 2009).

As expected, when transformed with the wild type Act-Cup reporter, alleles that had previously shown a temperature or cold sensitive growth phenotype also exhibited decreased growth on copper, consistent with inefficient splicing. Temperature sensitive *brr2-C520R* exhibited reduced growth at 30°C and 37°C, while cold sensitive *brr2-R1107A, brr2-R1107L*, and *brr2-1* showed reduced growth at 25°C, 30°C, and 37°C (Figure 3D column 1). As compared to observing *brr2-RP* growth without a reporter and on rich media, the expanded detrimental temperature ranges observed in this assay demonstrate its sensitivity.

The mutated reporter data are more complex. Reporter effects vary between *brr2-RP* strains in both the direction and magnitude of their effect on growth (Figure 3D). Every single *brr2-RP* strain exhibited better growth (yellow coloring), indicative of error-prone splicing, in the presence of at least one Act-Cup reporter, though in some cases the effect was very slight. Furthermore, every single *brr2-RP* strain exhibited worse growth (blue coloring), indicative of inefficient or hyperaccurate splicing, in the presence of at least one Act-Cup reporter (Figure 3D). This pattern is unique to *brr2-RP*, as *brr2-1* showed only reduced growth and the growth of *brr2-R1107M* was identical to that of *BRR2* for all reporters tested, consistent with its behavior in previous assays (Figure 3D).

**Figure 3.**
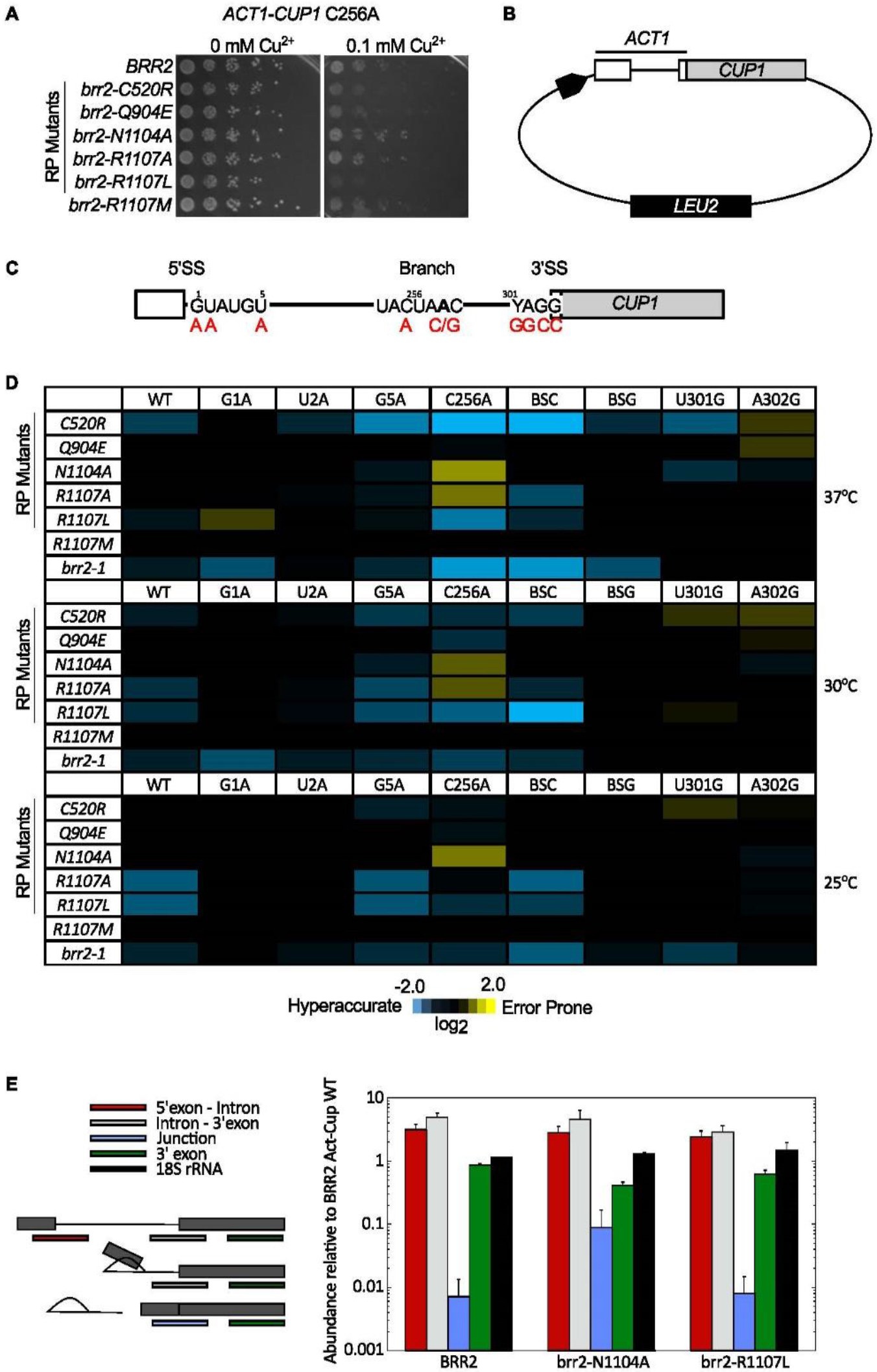
Brr2 is a splicing fidelity factor. A. Example of Act-Cup *brr2-RP* reporter assay. Indicated strains were spotted on minimal media containing CuSO_4_ and monitored for growth at 30°C. The left panel shows growth of *brr2-RP* yeast carrying a C256A Act-Cup reporter in the presence of 0 mM CuSO_4_, the right at 0.1 mM CuSO_4_. *brr2-N1104A* exhibits improved growth as compared to *BRR2*, while *brr2-R1107A* and *brr2-R1107L* growth is impaired. B. Schematic of Act-Cup reporter. The 5′ exon and intron of *ACT1* have been fused to the coding sequence of *CUP1*. Expression is driven by the GPD promoter. *LEU2* is included in the reporter plasmid as a selectable marker. C. Schematic of Act1-Cup1 pre-mRNA. Splice-site mutations made to test how *brr2-RP* impact splicing fidelity are in red. Heat map showing results of Act-Cup assays. Values shown were calculated by log_2_([Cu_max_ Prp8-RP]/[Cu_max_ Prp8-WT]) (Mayerle and Guthrie 2016). Blue indicates reduced growth in copper-containing media (hyperaccuracy) of *brr2-RP* relative to wild type with the same reporter, while yellow indicates improved growth (error-prone splicing). E. Results of a primer extension assay monitoring the abundance of different RNA species. Primers spanning the 5′exon intron junction (red) report on pre-mRNA abundance. Primers spanning the intron 3′exon junction (gray) report on pre-mRNA and lariat intermediate abundance. Primers spanning the 5′exon 3′exon junction (blue) report on spliced mRNA abundance. Primers within the 3′ exon (green) report on pre-mRNA, lariat intermediate, and mRNA levels. 18S rRNA was included as a control. The increased growth of *brr2-N1104A* carrying the C256A reporter is due to increased levels of Act-Cup mRNA. Data are the average of 3 biological measurements, error bars are SEM. Primer sequences provided in (MacRae et al. 2018).

**Figure 4.**
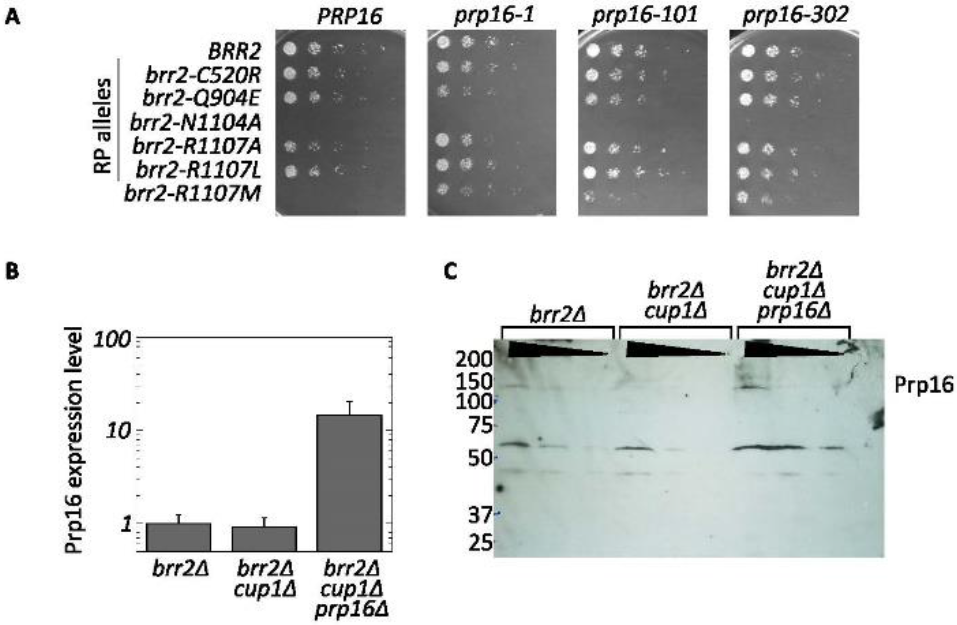
*brr2-RP* are sensitive to Prp16 abundance and ATPase activity. A. Growth assay. *brr2 prp16* double mutant yeast containing the indicated alleles were spotted onto rich media and allowed to grow at 30°C. In these yeast, the genomic copies of *BRR2* and *PRP16* have been knocked out and Brr2 and Prp16 are expressed from plasmids. In this strain background *brr2-N1104A* is lethal in combination with *PRP16* and all *prp16* alleles. *brr2-R1107M* is similarly lethal when combined with *PRP16* in this strain background, however, lethality can be rescued by ATPase deficient *prp16* alleles. B. RT-qPCR comparing *PRP16* transcript abundance (Primers: Prp16F 5′ AGGGCAAGAGACATTCGAGA 3′, Prp16R 5′ TGCTTGATGAGCAAATCCTG 3′). The strain used in panel A has increased *PRP16* mRNA abundance, likely due to its expression from a plasmid, not the genomic copy of *PRP16*. C. Western blot probed with polyclonal Prp16 antibody(Burgess and Guthrie 1993a) showing elevated Prp16 protein expression in the strain used in panel A. A nonspecific band at 60 KDa serves as a loading control.

The most interesting patterns occur in the presence of the C256A branch site mutant Act-Cup reporter. This reporter negatively affects the growth of *brr2-C520R*, *brr2-R1107L*, *brr2-R1107M*, and *brr2-1* at all temperatures consistent with inefficient or hyperaccurate splicing. At the same time, *brr2-N1104A* and *brr2-R1107A* growth is improved, indicative of error-prone splicing (Figure 3D, column 5). *brr2-Q904E* and *brr2-R1107M* grew as wild type *BRR2*. qRT-PCR analysis of the relative abundances of Act-Cup reporter pre-mRNA, lariat-intermediate, and mRNA are consistent with these growth data (Figure 3E). These data show that, at least in the case of *brr2-RP*, splicing fidelity is <u>not</u> directly linked to helicase activity or ATPase activity (Figure 1C, Figure 3D). Together our Act-Cup reporter data clearly implicate Brr2 in splicing fidelity, however the mechanism underlying its effects remains unclear.

### Prp16 ATPase activity and expression levels impact Brr2 function

While Brr2 has been shown to interact physically and genetically with Prp16(Fromont-Racine et al. 1997; Dix et al. 1998; van Nues and Beggs 2001; Liu et al. 2006; Hahn et al. 2012; Cordin et al. 2014; Absmeier et al. 2015; Galej et al. 2016), how these two proteins influence each other is unknown. Because both Brr2 and Prp16 are involved in proofreading the pre-mRNA branch site (Figure 3) (Burgess and Guthrie 1993a), and Prp16 ATPase activity has been directly linked to its proofreading activity(Burgess and Guthrie 1993b), we decided to determine if Prp16-ATPase deficient alleles genetically interact with *brr2-RP*. We created double mutant strains in which the genomic copies of both *BRR2* and *PRP16* are deleted and Brr2 and Prp16 are expressed from plasmids. We then shuffled in plasmids containing *BRR2* or *brr2-RP* and *PRP16* or a *prp16* ATPase deficient allele. Finally we plated these strains on rich media and assessed their growth at 30°C.

Unexpectedly, we observed synthetic lethality with *brr2-N1104A* and *brr2-R1107M* in combination with wild type *PRP16* when both proteins were expressed from plasmids (Figure 4A). Both Brr2 and Prp16 are expressed from low-copy plasmids in this strain, and both are under the control of their native promoters. However neither *BRR2* nor *PRP16* are expressed in their full genetic and chromosomal context. We therefore speculated that the synthetic lethality observed may be due to fact that Prp16 is overexpressed in this strain background. We assessed Prp16 expression at the mRNA level by RT-qPCR, and at the protein level by western blot with a polyclonal anti-Prp16 antibody. At both the mRNA and protein levels, Prp16 is overexpressed in the *brr2 prp16* double mutant strain as compared to strains where only Brr2 is expressed from a plasmid (Figure 4B,C). More intriguingly, *brr2-R1107M PRP16* synthetic lethality is rescued by ATPase deficient alleles of *prp16*, though these stains are clearly still sick (Figure 4A). All other combinations of *brr2-RP* and *prp16* alleles showed no genetic interactions. The interactions between *brr2-N1104A* and *brr2-R1107M* and *PRP16* and its ATPase deficient alleles do not correlate with any other phenotype tested, suggesting that erroneous interactions with Prp16 may be another way that *brr2-RP* negatively affects splicing. Because *brr2-R1107M PRP16* lethality is rescued by *prp16* ATPase alleles, these data suggest a functional link between Brr2 and Prp16 ATPase activity. However, more work is required to fully understand the mechanism underlying these genetic observations.

## Conclusion

Together these data implicate Brr2 in splicing fidelity, and highlight a previously unknown connection between Brr2 and Prp16 ATPase activity. They further implicate error-prone splicing in *SNRNP200*-related RP pathogenesis. More study is needed to understand how Brr2 impacts splicing fidelity, how the activities of Brr2 and Prp16 are coordinated, and what the functional consequences of disrupting these Brr2 functions are in Retinitis Pigmentosa.

## Materials and Methods

### Strains and Plasmids

Strains and plasmids were constructed using standard cloning techniques(Guthrie and Fink 2004). Act-Cup reporter plasmids were described in (Lesser and Guthrie 1993). All plasmids referenced in this manuscript, including the Act-Cup reporter plasmids, are available from Addgene (www.addgene.org). Strain requests should be directed to Hiten Madhani.

| STRAIN | GENOTYPE |
| --- | --- |
| YMEG022 | *brr2::KAN his leu lys ura yCP50-BRR2* |
| YMEG010 | *brr2::KAN cup1::ura3-52 his leu lys ura ade yCP50-BRR2* |
| YMEG766 | *brr2::KAN prp16::LYS cup1::ura3-52 ade met leu his yCP50-BRR2 pRS316-PRP16* |

### Yeast growth assays

For competitive growth assays, saturated overnight yeast cultures were diluted to OD_600_ of 0.1 in rich media and allowed to grow until they reached mid-log phase. Then the cultures were serially diluted and spotted onto rich media. Plates were allowed to grow at the indicated temperatures and then photographed.

### RT-qPCR

In all cases, total RNA was isolated from mid-log yeast cultures and prepped using the hot phenol method, treated with DNase, and then phenol-chloroform extracted again. cDNA was created using dN9 primers and Superscript III Reverse Transcriptase using the manufacturers protocols. Methods and primers for RT-qPCR of Act-Cup reporter cDNA are described in (MacRae et al. 2018), methods and primers for RT-qPCR of endogenous intron-containing transcripts are described in (Mayerle and Guthrie 2016). Prp16 primers are noted in the legend of Figure 4.

### ACT-CUP reporter assays

The Act-Cup reporter system has been described previously (Lesser and Guthrie 1993). The experiments described in this manuscript were performed using the same protocol as described in (Mayerle and Guthrie 2016).

### Western blot

Western blot analysis of Prp16 protein abundance was performed using the antibodies and method described in (Burgess and Guthrie 1993a).

## Acknowledgements

We thank everyone who has provided input into the interpretation of these data, particularly John Abelson, Jean Beggs, Corina Maeder, Magda Konarska, Charles Query, Jon Staley, Joan Steitz, Hiten Madhani, Jordan Burke, Sara Espinosa, and the UCSC RNA Club.

